# Deceptive safety? The impact of costly pain avoidance on the modulation and extinction of visceral pain-related fear

**DOI:** 10.1101/2025.06.25.661529

**Authors:** Franziska Labrenz, Anne Kalenbach, Sigrid Elsenbruch, Adriane Icenhour

**Affiliations:** Department of Medical Psychology and Medical Sociology, Ruhr University Bochum, Bochum, Germany; Department of Affective Neuroscience, Ruhr University Bochum, Bochum, Germany; Department of Neurology, Center for Translational Neuro- and Behavioral Sciences, University Hospital Essen, University of Duisburg-Essen, Germany

**Author notes:** **Corresponding author**: Adriane Icenhour, PhD, Department of Affective Neuroscience, Ruhr University Bochum, Universitaetsstrasse 105, 44789 Bochum, Germany.

**Keywords:** avoidance, extinction, fear conditioning, gut-brain-axis, pain-related fear, visceral pain

## Abstract

Along the gut-brain axis, visceral pain demonstrably evokes emotional learning and memory processes shaping behavior in clinically relevant ways. Avoidance motivated by learned fear may constitute a major obstacle to treatment success in extinction-based interventions. However, the effects of avoidance on visceral pain-related fear extinction remain poorly understood. By implementing an ecologically valid experimental protocol, we investigated how costly avoidance affects the modulation and extinction of visceral pain-related fear.

Thirty-three healthy volunteers underwent conditioning with visual cues (conditioned stimuli; CS^+^,CS^−^) consistently followed by visceral pain or remaining unpaired. During avoidance, participants decided to avoid or receive pain upon confronting CS^+^. Avoidance decisions resulted in pain omission in some trials, while in others, participants experienced unpredictable pain. During extinction, CS were presented unpaired. CS valence, fear, and trial-by-trial decisions were analyzed.

Avoidance decisions depended on prior experiences, with the highest probability of avoidance following successful pain omission. Negative CS^+^ valence and fear remained elevated across avoidance and extinction. Learned fear and more avoidance decisions explained 57% variance in sustained CS^+^ fear.

Our findings indicate that avoidance, which provides short-term absence of pain even when followed by unpredictable pain, motivates its maintenance. However, it perpetuates pain-related fear and may impede extinction, with implications for persisting symptoms and therapeutic outcomes in chronic visceral pain.

## Introduction

The ability to flexibly respond to environmental challenges is fundamental for maintaining or restoring homeostasis and ultimately ensures survival. Impending bodily harm, in particular, carries a high threat value and motivates associative learning, promoting the formation of classically conditioned associations between the imminent threat and its predictive cues. These associations allow the organism to initiate a cascade of protective responses, including increased arousal and avoidance. According to the fear avoidance model (FAM), an influential framework for the pathophysiology of anxiety disorders as well as chronic pain, avoidance and safety-seeking behaviors are fueled by fear of symptoms and reinforced through operant learning mechanisms^1,2^. However, if excessive fear and its behavioral consequences persist, avoidance loses its adaptive function and may even trigger detrimental effects, rendering the formerly protective cascade maladaptive and initiating a vicious cycle that can culminate in the development and persistence of disease^2,3^.

Building on this model, recent research has emphasized not only the role of fear in driving avoidance but also the costs associated with persistent avoidance behavior itself. Particularly in the context of chronic pain, costly avoidance refers to a behavioral strategy of persistently avoiding situations, activities, or internal sensations related to the threat of symptoms, even though this avoidance results in significant negative personal, psychological, or functional consequences^1,4^. While avoidance behaviors promote a sense of predictability of aversive sensations^5^, motivating their preservation, the short-term feeling of control is frequently followed by the unexpected and therefore particularly burdensome re-occurrence of symptoms, as previously shown in the context of movement-related pain^6,7^. This may not only perpetuate fear^8,9^ but also undermine the effectiveness of cognitive behavioral therapy (CBT), particularly exposure-based interventions^7,10–12^ that rely on the principles of extinction learning ^3,13,14^.

Despite the significant relevance to both pathophysiology and therapy of chronic pain conditions, for a long time pain-related avoidance behavior and its underlying mechanisms have not received the attention they deserve in experimental research. Typically, experimental avoidance research has operationalized avoidance costs as the expense of something valuable, such as time or money^2,15^. More recent studies, however, have broadened this scope to also incorporate intangible costs. In the context of musculoskeletal pain, ecologically valid models introducing avoidance costs as increased effort in performing movements have been applied to investigate avoidance behaviors in modulating pain-related fear and their key role as mechanisms of action in treating chronic pain^16,17^. However, with these few exceptions, ecologically valid paradigms that take clinically relevant avoidance costs and the impact of prior experiences into account are still largely absent, which limits translation into patients’ clinical reality, particularly regarding interoceptive visceral pain as a key characteristic in disorders of gut-brain interaction, such as irritable bowel syndrome (IBS).

IBS is characterized by recurrent pain accompanied by alterations in bowel habits, which patients describe as particularly unpredictable and burdensome^18,19^. This results in various, yet often unsuccessful attempts to identify and avoid interoceptive or exteroceptive predictors of symptoms, ranging from certain foods or drinks and tight clothing to activities and social interactions, recently summarized as “trial-and-error nature of identifying triggers”^20^. These distinct features of avoidance, together with recurrent failures to avoid pain resulting in unpredictable symptoms^21^, constitute key factors to disability, absenteeism, and social withdrawal^20^. These long-term costs of avoidance may likely also contribute to frequent comorbidities with anxiety and depression in disorders of gut-brain interaction^22^.

Not only from a clinical, but also an experimental perspective, visceral pain may be particularly susceptible to fostering avoidance and safety behaviors. It is demonstrably more unpleasant and threat-inducing than somatic pain in both healthy individuals^23–25^ and patients with chronic visceral pain^26,27^. Associative learning paradigms involving visceral stimuli have been shown to elicit robust behavioral, psychophysiological, and neural pain-related responses^28–32^, and importantly, to distinctly shape safety-related effects in both health^33,34^ and chronic visceral pain^35,36^. In line with the abovementioned clinical observations, ambiguity or the lack of reliable predictability appears to play a particular role, contributing to hypervigilance and anticipatory anxiety in healthy individuals as well as patients with IBS^37,38^. Ultimately, learned visceral pain-related fear may particularly encourage avoidance responses, with potential implications for sustained fear and aberrant extinction learning, which yet remains to be elucidated.

Based on this previous work in the field of fear conditioning, applying an established clinically relevant visceral pain model, we herein implemented a novel ecologically valid experimental pain-related avoidance paradigm to elucidate behavioral and psychological underpinnings of visceral pain-related fear learning, avoidance, and extinction. Specifically, we aimed to test the hypothesis that (1) the magnitude of classically conditioned pain-related responses would predict subsequent avoidance decisions. By operationalizing avoidance costs as the exposure to the threatening experience of unpredictable pain, our paradigm further enabled us to test the hypothesis that (2) prior avoidance decisions and their consequences in terms of successful pain avoidance or unexpected pain as avoidance costs would impact subsequent avoidance decisions. Finally, we hypothesized that (3) individual trial-by-trial avoidance choices and consequences promote the persistence and impede the extinction of pain-related fear.

## Results

### Sample characterization

An overview of sociodemographic and psychological variables is provided in Table 1. Briefly, mean age and BMI were well within range, with men (mean age 27.61 ± 0.78 years) being marginally older than women (mean age 25.07 ± 0.82 years) in the current sample. Participants presented with overall few psychological symptoms and reported low chronic stress, consistent with stringent exclusion criteria. Visceral sensory, urge, and pain thresholds were in accordance with previous findings on visceral pain sensitivity in young healthy volunteers^39^. Men and women did not differ with respect to any psychological measures or thresholds.

**Table 1.** Sociodemographic and psychological characterization of the study sample.

|  | Mean [min - max] | SEM | *p* ♀ vs. ♂ |
| --- | --- | --- | --- |
| Age | 26.45 [19 - 33] | 0.60 | .032 |
| BMI (kg/m<sup>2</sup>) | 24.10 [18.01 - 30.48] | 0.54 | .779 |
| Sensory threshold | 11.06 [6 - 22] | 0.88 | .675 |
| Urge threshold | 20.06 [10 - 34] | 1.03 | .958 |
| Pain threshold | 32.26 [14 - 48] | 1.64 | .746 |
| Emotional distress (HADS) |  |  |  |
| Anxiety | 3.79 [0 - 11] | 0.47 | .363 |
| Depression | 1.79 [0 - 6] | 0.31 | .730 |
| Trait anxiety (STAI-T) | 32.91 [20 - 45] | 1.22 | .898 |
| Chronic stress load (TICS) | 13.64 [0 - 32] | 1.39 | .506 |
| Intolerance of Uncertainty (UI18) |  |  |  |
| Reduced ability to act | 11.33 [6 - 20] | 0.57 | .834 |
| Burden | 13.52 [6 - 24] | 0.79 | .923 |
| Vigilance | 15.82 [7 - 25] | 0.91 | .382 |
| Pain-related cognitions (PRSS) |  |  |  |
| Catastrophizing | 0.86 [0 - 4] | 0.15 | .086 |
| Coping | 3.58 [2 - 5] | 0.14 | .726 |
| Resilience (RS-13) | 72.88 [57 - 90] | 1.29 | .512 |
| Self-efficacy (SWE) | 33.03 [21 - 66] | 1.22 | .644 |

Comparisons between women and men revealed no sex differences in visceral sensitivity or any relevant psychological markers. Data are provided as mean (with minimum and maximum values in square brackets), and SEM and *p* values are derived from two sample t-tests comparing male and female participants. Abbreviations: BMI, body mass index; HADS, Hospital Anxiety and Depression Scale; PRSS, Pain-Related Self Statements; RS, Resilience Scale; STAI, State Trait Anxiety Inventory; SWE, Self-efficacy Expectation; TICS, Trier Inventory of Chronic Stress; UI, Uncertainty Intolerance.

### US-related measures across experimental phases

Ratings of US pain intensity (*F*_(1.91,61.04)_ = 8.31; *p* = .001; *η_p_*^2^ = .21), valence (*F*_(1.87,59.81)_ = 3.70; *p* = .033; *η_p_*^2^ = .10) and fear of US (*F*_(1.98,63.25)_ = 6.89; *p* = .002; *η_p_*^2^ = .18) revealed significant main effects of *time*. Post hoc tests showed significant increases from baseline to post-acquisition values for intensity *t*_(32)_ = 2.99; *p* = .015; *d* = 0.52), valence (*t*_(32)_ = 2.60; *p* = .042; *d* = 0.45) and fear of US (*t*_(32)_ = 3.03; *p* = .015; *d* = 0.53) (Figure 1). No changes from post-acquisition to post-avoidance were detectable for US-related measures of valence (*t*_(32)_ = 0.15; *p* > .999; *d* = 0.03), fear (*t*_(32)_ = 0.09; *p* > .999; *d* = 0.02) and intensity (*t*_(32)_ = 0.65; *p* > .999; *d* = 0.11). Exploratory correlational analyses revealed that the percentage of avoidance choices after acquisition showed no relation with either US intensity (*p* > .191), valence (*p* > .184), or fear of US (*p* > .068).

**Figure 1.**
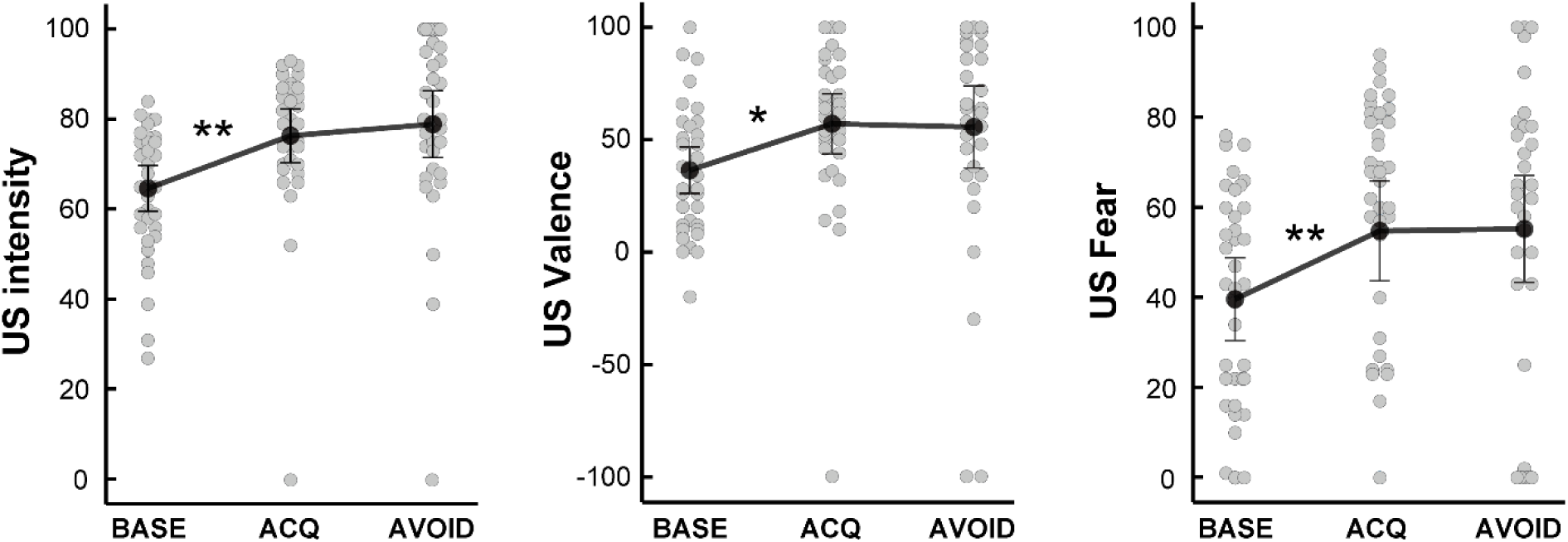
Ratings of perceived (A) intensity, (B) valence and (C) fear of visceral US assessed at baseline and after acquisition and avoidance phases. Data are provided as means and error bars indicate SEM. ** *p* < .01, * *p* < .05. Abbreviations: BASE, Baseline; ACQ, Acquisition, AVOID, Avoidance; US, unconditioned stimulus.

### Correlates of predictive cue properties across experimental phases

Results from RM-ANOVA of CS valence, fear and CS-US contingencies are provided across all experimental phases and findings from post hoc t-tests for the respective phase (acquisition, avoidance, extinction) are reported below. RM-ANOVA of CS valence revealed main effects of *CS type* (*F*_(1.57,48.69)_ = 17.31; *p* < .001; *η_p_*^2^ = .36) and *time* (*F*_(2.29,71.01)_ = 10.32; *p* < .001; *η_p_*^2^ = .25), as well as a *CS type* x *time* interaction (*F*_(4.17,129.21)_ = 9.65; *p* < .001; *η_p_*^2^ = .24). For fear of CS, analysis yielded main effects of *CS type* (*F*_(1,32,40.92)_ = 25.23; *p* < .001; *η_p_*^2^ = .45) and *time* (*F*_(2.61,80.92)_ = 15.31; *p* < .001; *η_p_*^2^ = .33), and a *CS type* x *time* interaction (*F*_(3.43,106.16)_ = 11.40; *p* < .001; *η_p_*^2^ = .27). Finally, analysis of CS-US contingency ratings indicated main effects of *CS type* (*F*_(1.67,51.85)_ = 120.06; *p* < .001; *η_p_*^2^ = .80) and *time* (*F*_(1.00,31.00)_ = 35.93; *p* < .001; *η_p_*^2^ = .54), as well as a *CS type* x *time* interaction (*F*_(1.42,43.99)_ = 37.35; *p* < .001; *η_p_*^2^ = .55).

### Post hoc tests for pain-related fear acquisition

Results of post hoc tests regarding CS valence and fear of CS as self-report correlates of learned emotional responses to predictive cues are depicted in Figure 2. For CS valence, post hoc tests demonstrated a significant increase of CS^+^ valence induced by pain-related acquisition (*t*_(32)_ = 7.64; *p* < .001; *d* = 1.33), whereas ratings of CS^−^ (*t*_(32)_ = 0.24; *p* > .999; *d* = 0.04) and CS^0^ (*t*_(32)_ = 1.12; *p* = .816; *d* = 0.19) remained stable. Accordingly, significant differences between CS^+^ and both, CS^−^ (*t*_(32)_ = 6.35; *p* <.001; *d* = 1.10) and CS^0^ (*t*_(32)_ = 4.49; *p* =.002; *d* = 0.78) were observable at the end of the acquisition phase, while valence ratings of CS^−^ and CS^0^ did also differ (*t*_(32)_ = 2.73; *p =* .030; *d* = 0.48) (Figure 2A).

**Figure 2.**
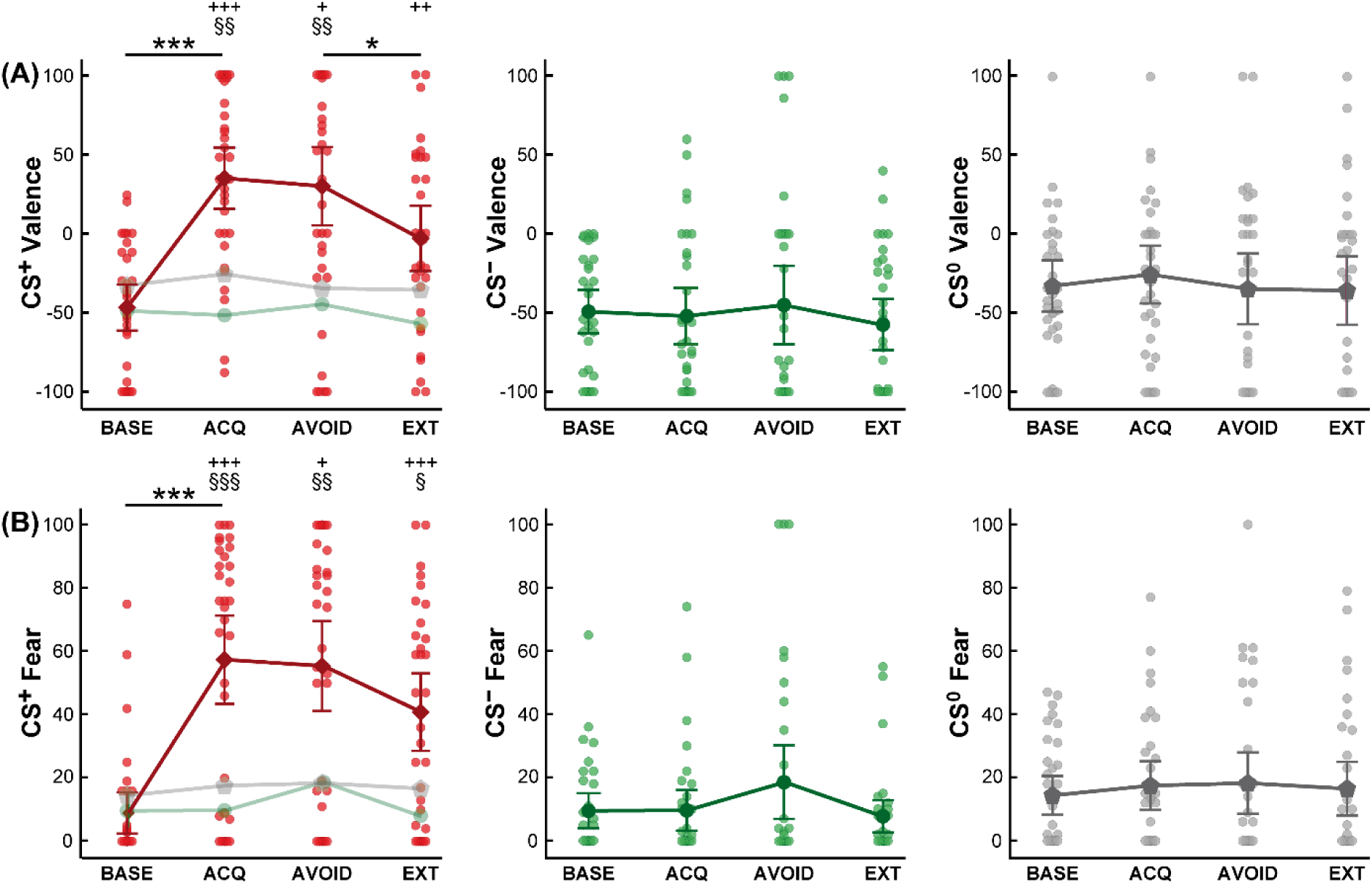
Ratings of (A) valence and (B) fear regarding CS^+^ (left panel, depicted in direct comparison to CS^−^ and CS^0^ indicated in light colors), CS^−^ (middle panel) and CS^0^ (right panel) across experimental phases. Data are provided as means and error bars indicate SEM. Significant differences for post hoc comparisons between experimental phases are indicated with *** *p* < .001 and * *p* < .05, comparisons between CS^+^ and CS^−^ within experimental phases with +++ *p* < .001, ++ *p* < .01 and + *p* < .05 as well as between CS^+^ and CS^0^ with §§§ *p* < .001, §§ *p* < .01 and § *p* < .05. Abbreviations: BASE, Baseline; ACQ, Acquisition, AVOID, Avoidance; CS, conditioned stimulus; EXT, Extinction.

For fear of CS, post hoc tests revealed a significant increase of CS^+^ fear induced by pain-related learning (*t*_(32)_ = 7.68; *p* <.001; *d* = 1.34), whereas fear ratings of CS^−^ (*t*_(32)_ = 0.08; *p* > .999; *d* = 0.01) and CS^0^ (*t*_(32)_ = 1.16; *p* =.771; *d* = 0.20) did not show any changes. After acquisition, CS^+^ fear ratings were significantly higher compared to CS^−^ (*t*_(32)_ = 6.88; *p* < .001; *d* = 1.20) as well as CS^0^ (*t*_(32)_ = 5.33; *p* < .001; *d* = 0.93), whereas CS^−^ and CS^0^ ratings showed no significant difference after correcting for multiple comparisons (*t*_(32)_ = 2.10; *p* = .132; *d* = 0.37) (Figure 2B).

Contingency ratings indicated clear awareness of CS-US associations following acquisition, with significantly higher ratings for CS^+^ (90.70 ± 2.63 %) when compared to CS^−^ (13.36 ± 4.13 %) (*t*_(32)_ = 13.31; *p* < .001; *d* = 2.32) and CS^0^ (10.73 ± 3.63 %) (*t*_(32)_ = 14.39; *p* < .001; *d* = 2.50), and no difference between CS^−^ and the control cue CS^0^ (*t*_(32)_ = 0.60; *p* > .999; *d* = 0.10).

### Post hoc tests for pain-related avoidance

All participants made an active decision to either avoid or receive pain when prompted (i.e., painful US were never delivered as a result of abstaining from or forfeiting a decision). On average, participants decided to avoid the US in 55.49 % of all avoidance trials, evenly distributed throughout the avoidance phase, depicted as individual percentages of avoidance decisions in Figure 3A. Across all trials, participants had a mean reaction time of 1.28 ± 0.10 s to reach their decision with no significant differences between times required for a decision to avoid or to receive the US, respectively (*p* = .470; Figure 3B, Table 2).

**Figure 3.**
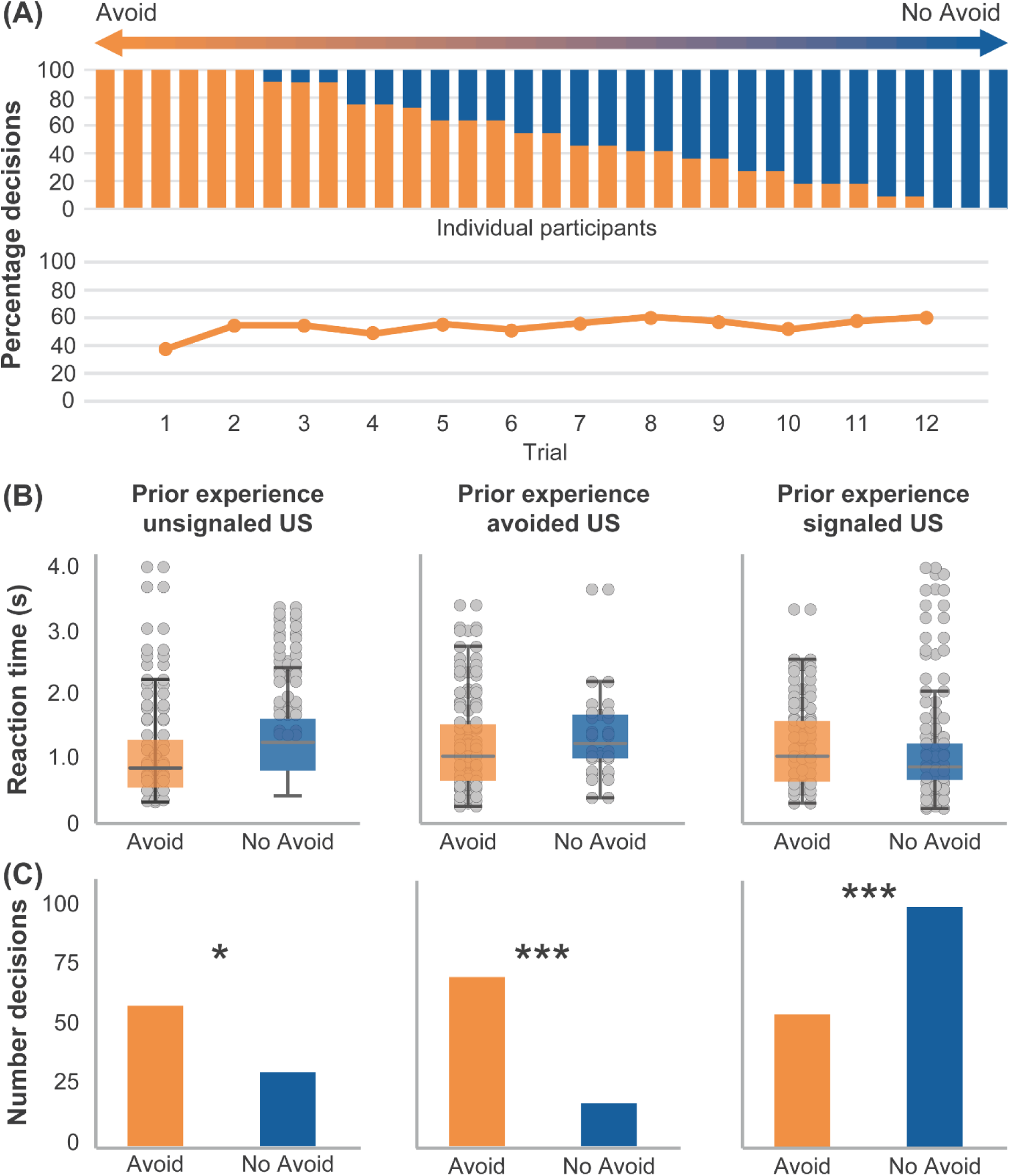
(A) Percentage of avoidance decisions in individual participants during avoidance (upper panel) and across trials (lower panel), (B) reaction times (in seconds) to reach a decision to avoid or receive the following US and (C) the total number of respective decisions. (B) and (C) were subdivided with respect to the pain-related experience preceding the decision, i.e. depicting (B) reaction times and (C) total number of decisions following a prior experience of an unpredicted unconditioned stimulus (US, left panel), following successful avoidance (middle panel) or decisions following the prior experience of a predicted US in the preceding trial (right panel). Data on reaction times in seconds are given as mean ± SEM. *** *p* < .001, * *p* < .05, based on results of Fisher’s Exact Tests. Statistical details are provided in Table 2.

**Table 2.**
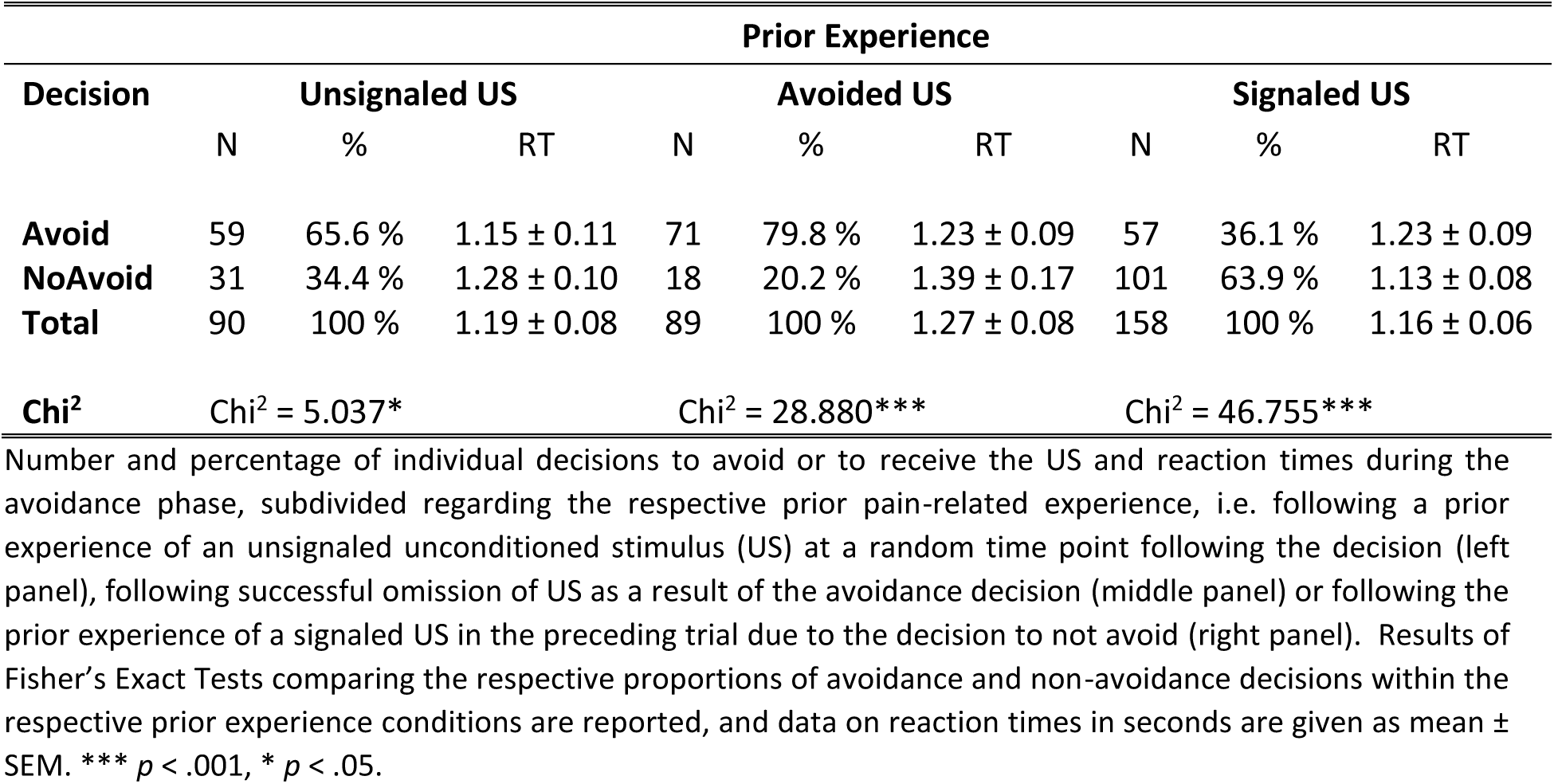
Decisions and reaction times subdivided regarding the respective prior experience during the avoidance phase.

A generalized linear mixed model analysis testing for trial-by-trial effects of prior pain-related experiences on subsequent decisions yielded a significant fixed effect of *prior experience* (unsignaled US: *F*_(1;323)_ = 125.29, *p* < .001; signaled US: *F*_(1;323)_ = 316.68, *p* < .001; avoided US: *F*_(1;323)_ = 140.11, *p* < .001) but no effect of *trial* (*F*_(10,323)_ = 0.47, *p* = .908) (Bayes = 1488.49). Effects were attributable to significant differences in proportions of decisions to avoid or not to avoid pain depending on the prior pain-related experience. Results of Fisher’s Exact Test revealed that when participants decided to not avoid and hence received a signaled, hence expected, US, they were more likely to make a subsequent choice of non-avoidance. On the other hand, the likelihood of choosing avoidance was highest following trials in which avoidance choices led to an omission of US. Statistical details from Fisher’s Exact Test are provided in Table 2, and Figure 3C depicts decisions during avoidance separately for the respective US experience during the preceding trial.

A generalized linear model for trial-by-trial analyses testing effects of trial and type of decision on reaction times yielded no significant main effects or interactions (all *p* > .189).

Analyses of CS valence ratings revealed that CS^+^ remained significantly more aversive compared to both, CS^−^ (*t*_(32)_ = 5.27; *p* =.015; *d* = 0.92) and CS^0^ (*t*_(32)_ = 4.49; *p* = .008; *d* = 0.78), and no difference between CS^−^ and CS^0^ (*t*_(32)_ = ; *p* > .999; *d* = 0.16) after the avoidance phase was detected (Figure 2A). Likewise, CS^+^ remained significantly more fear-inducing than CS^−^ (*t*_(32)_ = 5.32; *p* = .036; *d* = 0.91) and CS^0^ (*t*_(32)_ = 5.24; *p* = .003; *d* = 0.90), and no difference was observed between CS^−^ and CS^0^ (*t*_(32)_ = 0.60; *p* > .999; *d* = 0.01) (Figure 2B).

Hierarchical stepwise regression analyses were further conducted to complement findings from ANOVA, aiming to identify predictors of CS^+^ fear and valence following avoidance. Findings are summarized in Table 3. Briefly, CS^+^ valence rated after acquisition was identified as the strongest coefficient significantly explaining variance in CS^+^ valence at the end of avoidance. In contrast, neither fear nor the percentage of individual avoidance decisions was included in the model. Trait anxiety was further found to be a significant predictor, together accounting for 49% of the variance in CS^+^ valence ratings after the avoidance phase. For CS^+^ fear, on the other hand, learned fear of CS^+^ and avoidance decisions, but not valence, were included in the model as significant predictors. Together with trait anxiety and a reduced ability to act due to intolerance of uncertainty (UI18-A) as psychological variables, the model explained almost 70% of the variance in CS^+^ fear ratings at the conclusion of the avoidance phase. Regression analysis evaluating whether classically conditioned pain-predictive cue properties of valence and fear or psychological traits could explain variance in the percentage of subsequent avoidance decisions yielded no significant findings (all *p* > .126).

**Table 3.** Stepwise hierarchical regression analyses to predict emotional responses after avoidance.

| **CS<sup>+</sup> Valence** |  |  |  |  |  |  |
| --- | --- | --- | --- | --- | --- | --- |
| Variable | Stage 1 |  |  | Stage 2 |  |  |
|  | Behavioral and self-report predictors |  |  | Psychological predictors |  |  |
| | $\beta$ | $t$ | $p$ | $\beta$ | $t$ | $p$ |
| ACQ: CS<sup>+</sup> Valence | .632 | 4.54 | **<.001** | .588 | 4.45 | **<.001** |
| ACQ: CS<sup>+</sup> Fear | -.349 | 1.30 | .204 | -.191 | 0.71 | .485 |
| Percentage Avoided | .228 | 1.69 | .102 | .160 | 1.19 | .245 |
| Trait Anxiety |  |  |  | -.301 | 2.28 | **.030** |
| $R^2$ | | .399 | | | .488 | |
| Change in $R^2$ | | .399 | | | .088 | |
| $F$ | | 20.61 | | | 14.29 | |
| $p$ | | **<.001** | | | **<.001** | |
| **CS<sup>+</sup> Fear** |  |  |  |  |  |  |
| ACQ: CS<sup>+</sup> Valence | .231 | 0.93 | .360 | .006 | 0.03 | .980 |
| ACQ: CS<sup>+</sup> Fear | .705 | 5.54 | **<.001** | .671 | 6.38 | **<.001** |
| Percentage Avoided | .272 | 2.27 | **.031** | .232 | 2.12 | **.043** |
| Trait Anxiety |  |  |  | -.311 | 2.45 | **.021** |
| UI18 – Reduced Ability to Act |  |  |  | .398 | 3.23 | **.003** |
| $R^2$ | | .571 | | | .694 | |
| Change in $R^2$ | | .571 | | | .122 | |
| $F$ | | 35.85 | | | 15.87 | |
| $p$ | | **<.001** | | | **<.001** | |

Abbreviations: ACQ, Acquisition; CS, conditioned stimulus; UI, Uncertainty Intolerance.

**Table 4.** Stepwise hierarchical regression analyses to predict emotional responses after extinction.

| ***CS<sup>+</sup> Valence*** |  |  |  |  |  |  |
| --- | --- | --- | --- | --- | --- | --- |
| **Variable** | **Stage 1** |  |  | **Stage 2** |  |  |
|  | **Behavioral and self-report predictors** |  |  | **Psychological predictors** |  |  |
| | $\beta$ | $t$ | $p$ | $\beta$ | $t$ | $p$ |
| ACQ: CS<sup>+</sup> Valence | .082 | 0.43 | .669 |  |  |  |
| ACQ: CS<sup>+</sup> Fear | -.018 | 0.11 | .916 |  |  |  |
| AVOID: CS<sup>+</sup> Valence | .581 | 3.98 | **<.001** |  |  |  |
| AVOID: CS<sup>+</sup> Fear | .084 | 0.40 | .690 |  |  |  |
| Percentage Avoided | .176 | 1.17 | .251 |  |  |  |
| ∅ | | | | | | |
| $R^2$ | | .338 | | | | |
| Change in $R^2$ | | .338 | | | | |
| $F$ | | 15.84 | | | | |
| $p$ | | **<.001** | | | | |
| ***CS<sup>+</sup> Fear*** |  |  |  |  |  |  |
| ACQ: CS<sup>+</sup> Valence | -.052 | 0.25 | .804 |  |  |  |
| ACQ: CS<sup>+</sup> Fear | -.025 | 0.12 | .909 |  |  |  |
| AVOID: CS<sup>+</sup> Valence | .127 | 0.58 | .567 |  |  |  |
| AVOID: CS<sup>+</sup> Fear | .519 | 3.38 | **.002** |  |  |  |
| Percentage Avoided | .038 | 0.23 | .821 |  |  |  |
| ∅ | | | | | | |
| $R^2$ | | .270 | | | | |
| Change in $R^2$ | | .270 | | | | |
| $F$ | | 11.45 | | | | |
| $p$ | | **.002** | | | | |

Abbreviations: ACQ, Acquisition phase; AVOID, Avoidance phase; CS, conditioned stimulus.

### Post hoc tests for pain-related extinction

For CS valence ratings, post hoc tests showed that while CS^+^ was rated as significantly less unpleasant after the extinction phase when compared to the avoidance phase (*t*_(32)_ = 3.32; *p* =.036; *d* = 0.59), a significant difference when compared to CS^−^ was maintained (*t*_(32)_ = 4.32; *p* =.003; *d* = 0.76) (Figure 2A). No significant difference between avoidance and extinction was found for CS^−^ (*t*_(32)_ = 1.09; *p* > .999; *d* = 0.19) and CS^0^ (*t*_(32)_ = 0.13; *p* > .999; *d* = 0.02). The comparison between CS^+^ and CS^0^ (*t*_(32)_ = 2.47; *p* = .114; *d* = 0.44) and between CS^−^ and CS^0^ (*t*_(32)_ = 2.45; *p* =.012; *d* = 0.43) after extinction also yielded no significant difference.

Likewise, post hoc tests revealed that CS^+^ was still rated as significantly more fear-inducing compared to both, CS^−^ (*t*_(32)_ = 5.52; *p* <.001; *d* = 0.98) and CS^0^ (*t*_(32)_ = 3.84; *p* = .010; *d* = 0.68) after the extinction phase (Figure 2B). While ratings of fear decreased after extinction compared to levels after avoidance for all CS, these effects failed to reach statistical significance after Bonferroni-correction (CS^+^ (*t*_(32)_ = 2.31; *p* = .168; *d* = 0.41), CS^−^ (*t*_(32)_ = 1.85; *p* = .219; *d* = 0.33, CS^0^ (*t*_(32)_ = 0.43; *p* > .999; *d* = 0.08)).

To further substantiate findings from ANOVA, hierarchical stepwise regression analyses were conducted. For CS^+^ valence after extinction, a significant model was identified with CS^+^ valence after avoidance as the strongest predictor, accounting for 34% of variance. In contrast, measures acquired after acquisition, fear, percentage of avoidance choices, or psychological traits provided no additional predictive value to the model. Regarding CS^+^ fear after extinction, a model involving only fear ratings after avoidance as a predictor was identified, explaining 27% variance.

CS-US contingency ratings acquired after the extinction phase revealed that while differences between CSs were less pronounced and ratings for CS^+^-US contingencies were lower than after the acquisition phase (28.06 ± 6.23 %; *t*_(32)_ = 8.19; *p* < .001; *d* = 1.45), significance was not reached when comparing CS^+^ to CS^−^ (10.75 ± 3.42 %) or to CS^0^ contingency ratings (13.16 ± 4.09 %) after Bonferroni-correction (both *p* > .072). No significant differences in CS-US contingency were observed between acquisition and extinction for CS^−^ (*t*_(32)_ = 0.59; *p* > .999; *d* = 0.11) and CS^0^ (*t*_(32)_ = 0.45; *p* > .999; *d* = 0.08).

## Discussion

In the face of acute threat, avoidance constitutes an integral part of a highly adaptive behavioral repertoire aiming at self-protection and survival of the organism. Excessive and persistent avoidance, however, is a hallmark characteristic not only of anxiety disorders^40^, but also chronic pain^1^, and can be highly debilitating. Available treatment options, such as exposure-based treatments, aim at extinguishing fear of pain, which is a core incentive of avoidance and its detrimental effects, yet behavioral avoidance patterns often persist during and after treatment, even when accompanied by meaningful costs^41,42^. This can severely compromise the efficacy of exposure and hamper long-term success of these therapeutic approaches, which are particularly promising for patients suffering from chronic pain conditions in which fear plays a particular role, such as in disorders of bidirectional gut-brain interactions^19,43^. Despite this relevance for the pathophysiology and therapy of chronic visceral pain, ecologically and clinically valid research approaches addressing the impact of costly avoidance on inhibitory learning have thus far been uncharted territory, even in healthy samples. To our knowledge, the current study provides first evidence on the impact of costly avoidance on the modulation and extinction of visceral pain-related fear.

Confirming successful fear acquisition, negative valence, and fear of the pain-predictive cue along with widely accurate contingency ratings at the end of the acquisition phase, were well in line with our earlier observations in the context of differential visceral pain-related fear learning in both health and disease^23,24,28–32,34,36,44^. Irrespective of these consistent and robust fear responses, a substantial interindividual variability in decisions to avoid or receive visceral pain was observed throughout the subsequent avoidance phase. Specifically, subjects either persistently avoided, irrespective of experiencing costs in terms of pain that was not reliably predictable but applied at a random time point, changed their decision patterns, or made an active decision to receive each pain stimulus despite the possibility to omit it. This observation supports recent experimental findings identifying subgroups of healthy individuals characterized by pain approach and pain avoidance^45^, consistent with our observation of a substantial group of non-avoiders. On the other hand, the avoidance endurance model conceptualizes different patterns of response repertoires to pain and their impact on pain experiences and disability^46^. According to this, not only persistent avoidance but also endurance-related responses may constitute pathophysiological mechanisms contributing to chronicity, disability, and poor treatment outcome. This would not only have clear implications for more refined and personalized therapeutic approaches for patients with chronic, including visceral, pain, but also calls for more translational research to identify relevant modulators, including, but not limited to, pain attitudes, tendencies to catastrophize, or sensation seeking^45,46^.

In the current analysis, no psychological trait assessed was associated with these individual decision patterns, which must be interpreted in light of the young, healthy sample with overall low scores and variance in these clinically relevant trait markers. Instead, participants appeared to be mainly motivated by the prior pain-related experience. Specifically, individuals who decided to confront themselves with the US and hence instantly received visceral pain were more likely to repeat than to change their decision in the subsequent trial. Conversely, the likelihood to perpetuate avoidance was highest after successful pain omission, but, importantly, also elevated after the aversive experience of not being able to reliably predict visceral pain. Unlike hypothesized, conditioned fear did not predict avoidance patterns; instead, the proportion of avoidance decisions and the intensity of learned fear predicted its persistence. This indicates that, at least in young healthy individuals, fear is not a direct trigger of active avoidance decisions. Yet, avoidance and consequences of these decisions appear to fuel pain-related fear, eventually spiraling into a vicious circle. These findings complement recent experimental data in the context of musculoskeletal pain, supporting the persistence or even increase of perceived threat and fear when individuals engage in avoidance behavior^6,9,47,48^. Several explanations may account for this phenomenon, with the most recognized one being referred to as “behavior as information”^49,50^. Specifically, individuals may interpret their defensive behaviors as validation of an imminent threat, using their own avoidance responses to infer the presence of danger. This tendency appears to be especially pronounced in clinical populations^50,51^. Moreover, avoidance responses herein came with the cost of experiencing unexpected pain, which may also account for the association between avoidance and fear.

Our findings are intriguing both conceptually and clinically. Individuals who actively choose to confront themselves with pain could thereby gain control over their symptoms. This behavioral pattern aligns with the principles of the fear avoidance model and is considered a vital skill in cognitive-behavioral interventions for chronic visceral pain^1,52^. Albeit speculative, these individuals might be less susceptible to the “deceptive safety” that avoidance provides. Instead, they opt for pain experiences that they can prepare for and manage effectively. As a result, these coping resources could attenuate the connection between negative psychological factors, avoidance behaviors, and pain-related disability, which has significant implications for therapeutic interventions.

In line with the fear-avoidance model, avoidance, particularly if perceived as effective, may provide short-term absence of pain as a form of deceptive safety. This temporary state can be rewarding, encouraging the continuation of these behavioral patterns, yet conserving pain-related fear. Our data extend this perspective, documenting the perseveration of avoidance even when the decision results in a loss of predictability induced by an unexpected pain experience.

Ambiguity and uncertainty reportedly increase fear^53,54^ and could, in the context of pain-related associative learning, even contribute to central pain amplification^28,55^. Our data on pain perception demonstrating increased intensity, unpleasantness, and fear of pain after pain-related fear acquisition, which remained at stable, elevated levels during avoidance, lend support for hyperalgesic effects. Albeit no evident associations with avoidance decisions, this suggests that repeated exposure to pain or pain-related learning may have influenced both the sensory and emotional aspects of pain perception. Importantly, while findings from our healthy sample preclude directly transferable implications to clinical populations, the relative lack of basic knowledge regarding normal pain functioning must be considered. Therefore, data from healthy individuals can provide valuable insights to detect such risk factors or to date unidentified at-risk groups in research on the transition from acute to chronic pain. Ultimately, the observed phenomena may accumulate to persistent and maladaptive avoidance patterns in the transition from bothersome pain attacks and increasing, yet often unsuccessful, efforts to predict and control them to chronicity in visceral pain conditions^18,20,21^.

Excessive avoidance behaviors constitute transdiagnostic symptoms of anxiety disorders as well as chronic visceral pain. Although fear extinction as a fundamental mechanism underlying exposure-based treatment approaches^7,56,57^ has proven efficacious in various chronic pain conditions, pain-related fear may persist or return, hampering long-term treatment success^43,58^. Our results from a non-clinical sample showed that after extinction, during which no visceral pain stimulus was applied and no decisions were available, participants overestimated cue and pain contingencies, and negative valence and fear of the pain-predictive cue remained elevated, in line with the assumption that safety behaviors preserve threat beliefs^59^. While one possible explanation for the incomplete extinction of pain-related negative valence and fear could be an insufficient number of extinction trials, our previous studies using the same or even fewer trials during extinction did not yield similar persistence effects^24,28,31^. Instead, increased negative valence and fear, as assessed after the avoidance phase, were found to predict sustained negative emotional responses after extinction. In other words, sustained negative valence and fear after having the possibility to avoid, yet irrespective of individual avoidance decisions, may impact the persistence of pain-related fear even following extinction training. Further phenomena underlying the sustained responses may reflect impaired reconsolidation of extinction learning or avoidance-based disruption of fear memory updating, as two previously discussed mechanisms of action by which avoidance interferes with extinction. Prior work has shown that extinction forms a new inhibitory trace rather than erasing the original memory^60^, rendering fear particularly susceptible to renewal, reinstatement, or spontaneous recovery, especially when contextual or cognitive factors interfere with extinction consolidation^61^. The opportunity to avoid may thereby interfere with reconsolidation or updating of fear memories and compromise opportunities for corrective learning^62^. Reducing possibilities to avoid might therefore be a prerequisite for successful fear extinction and the long-term efficacy of exposure-based treatment approaches^19^. Future studies should focus on how to directly modify avoidance patterns, such as through the devaluation of the unconditioned stimulus (US) and cost-benefit analyses comparing approach versus avoidance strategies^63^. Understanding the neurobiology of pain-related fear avoidance could also open new avenues for behavioral, cognitive, or neural interventions.

Of note, some aspects of the current experimental approach may limit causal inferences and therefore deserve further consideration to inspire future translational approaches in pain-related avoidance research. Firstly, the active choice to either avoid pain or confront it when faced with a pain-predictive signal was provided in the majority, but not in all trials. From a clinical perspective, this circumstance may reflect the reality of patients suffering from disturbed gut-brain communication, where active avoidance might be available in many situations, yet is not always an option. In light of widely lacking direct evidence, one can only speculate whether, in chronic visceral pain conditions, an inability to avoid could promote more passive forms of safety behaviors or maladaptive coping strategies, resulting in the persistence of fear, a hypothesis worth investigating. Taking the learning theory perspective, one may conversely argue that these trials during the avoidance phase involving contingent CS^+^-US associations without the opportunity to avoid pain, resembled a partial reinforcement and may have distinctly maintained or even strengthened the pain-related memory trace. While cue-pain contingency ratings might support this notion, other observations are less in favor of this interpretation. For instance, if persistent negative emotional responses were predominantly driven by higher reinforcement rates, one could expect the highest fear and valence ratings among those individuals who consistently chose to confront pain. This would increase the number of cue-pain associations and strengthen the emotional memory related to pain. However, the observation that higher fear levels were predicted by a greater proportion of avoided pain experiences indicates the opposite. These patterns lend further support that perceived control regarding pain symptoms may play a more critical role in reducing pain-related fear than simply avoiding pain. A systematic assessment of perceived control and subjectively experienced costs is therefore a crucial extension in future avoidance research in the field of chronic visceral pain.

On a related note, successful avoidance, and consequently the omission of pain, was identified as the strongest predictor of maintaining this decision pattern. However, while the costs associated with avoidance seemed to influence decision changes to some extent, evidenced by a reduced proportion of avoidance choices after experiencing unexpected pain, certain participants displayed a rigid pattern of avoidance choices, even following repeated failures to avoid pain. Possibly, to these individuals, the costs may have been too subtle or infrequent to override the overall benefit of avoidance, highlighting the importance of the motivational context^64^ and underscoring the need for an assessment of subjective costs and values of avoidance. Current endeavors to optimize personalized treatments encourage a balanced approach to avoidance in therapeutic interventions^19,40,52,65^. Identifying relevant factors that predict beneficial or harmful effects of tolerating avoidance behaviors in exposure-based treatments for chronic pain presents a promising future direction. This would meet the articulated need for advanced diagnostic validity in translational research^5^ and could inspire targeted approaches taking the individual motivation and consequences of costly avoidance into account.

Secondly, causal inferences could be strengthened by including an appropriate control group, as previously accomplished in the field of musculoskeletal pain^9,47,48,66–68^. However, our experimental design allowed for the identification of individual learning and avoidance trajectories, focusing specifically on the role of individual learning histories and decisions, which inherently require participant agency. While this could potentially limit the comparability to a control group receiving the same amount of pain stimuli without considering their own prior behavior, establishing an adequate yoked control group appears a promising future direction in the field.

Thirdly, it is worth noting that our design relied on retrospective ratings obtained after the experimental blocks, which may be subject to recall bias and reduced temporal resolution. This approach was chosen to minimize cognitive load and avoid interrupting the associative learning process during the task induced by trial-by-trial assessments; however, it also poses a limitation and calls for the incorporation of online measures to enhance ecological and temporal validity in future studies.

Ultimately, further investigation into the mechanisms underlying visceral pain-related avoidance and its effects on sustained pain-related fear and resistance to extinction could provide valuable insights. These may hold significant implications for developing and refining tailored treatment options for patients suffering from chronic visceral pain related to disorders of the gut-brain axis.

## Materials and Methods

### Participants

The sample size was estimated in an a priori power analysis using the software G*Power (version 3.1.9.4^69^). Based on observations from previous fear conditioning studies with visual cues as conditioned and interoceptive visceral pain as unconditioned stimuli in healthy volunteers^24,28,30–33^, we assumed an average effect size of d = 0.49 or f = 0.25, a correlation among repeated measures of r = .30 and a non-sphericity correction factor Ɛ of 0.8 using a significance level of α = 0.05 and a power of 1-β ≥ 0.95 for valence and fear ratings in the planned study, resulting in a minimum total sample of N = 32. Accounting for a possible drop-out or incomplete datasets, 38 healthy individuals (18 women, 20 men, sex assigned at birth) were recruited by local advertisement. The recruitment and screening process included a structured telephone interview to screen prospective participants for the following inclusion criteria: Age between 18 and 45 years, body mass index (BMI) ≥ 18 and ≤ 30 kg/m^2^, and no history of acute or chronic gastrointestinal, neurological or psychiatric conditions or chronic pain conditions. We further excluded participants whose health status would necessitate the regular use of prescribed or over-the-counter medication, except hormonal contraceptives, thyroid medications, or occasional use of over-the-counter allergy or pain medication. If individuals were eligible, they were invited for a personal interview during which they received standardized study-related information and gave informed written consent. Participants were informed that, during the study, they would see visual stimuli and experience interoceptive visceral pain stimulation, but no information regarding experimental phases, contingency changes, the possibility to avoid pain or relations between visual cues and visceral stimulation was disclosed.

Current symptoms of emotional distress, particularly anxiety and depression, were screened using the German version of the Hospital Anxiety and Depression Scale (HADS^70^; α = .80) and participants scoring above a cutoff score of ≥ 8 were excluded. The frequency and severity of symptoms of the lower and upper gastrointestinal (GI) tract were assessed with a standardized in-house questionnaire^71^ and a score indicative of a GI condition (≥ 11) led to exclusion. Of note, the study was designed as a functional magnetic resonance imaging (fMRI) study. Only behavioral and self-report measures are reported herein, but the usual MRI-related exclusion criteria were applied, and all participants were right-handed, as confirmed with a validated questionnaire on motor asymmetries^72^. Structural brain abnormalities were excluded based on structural MRI on the study day. Only women using hormonal contraception were included in the study to reduce a putative confounding effect of fluctuations in sex steroid hormone concentrations across the menstrual cycle in female participants. This approach was chosen based on previous findings documenting effects of gonadal hormone concentrations and menstrual cycle phase on both the perception of visceral symptoms^73^ and emotional learning and memory processes^74,75^. Pregnancy was ruled out with a commercially available urinary test on the study day. A physical examination was conducted to exclude perianal tissue damage (i.e., fissures or painful hemorrhoids), which could interfere with the experimental procedures, particularly rectal balloon placement and distensions.

The study protocol was approved by the ethics committee of the Medical Faculty, University of Duisburg-Essen (protocol number 16–7226-BO) and followed the provisions of the Declaration of Helsinki. All participants gave written informed consent and received a financial compensation of 150 € each for their participation.

### Questionnaires

In addition to the HADS and the GI symptom questionnaire (see above), the following comprehensive questionnaire battery was provided to characterize the sample with respect to relevant psychological variables: The trait version of the State Trait Anxiety inventory (STAI-T^76^) to assess trait anxiety (sum scores between 20 and 80; α = .91), the Trier Inventory for the Assessment of Chronic Stress (TICS^77^) to measure chronic stress (sum scores 0-48; α = .87), the general scale for self-efficacy beliefs (SWE^78^) with sum scores between 10 and 40 (α = .80), the Uncertainty Intolerance Questionnaire (UI-18^79^) as a measure of trait intolerance to uncertainty (scores ranging from 18 to 90; α = .90), the Pain-Related Self Statement Scale (PRSS^80^) to measure pain-related cognitions in terms of maladaptive pain catastrophizing and adaptive pain coping (sum score range 0-45, respectively; α = .90) and the Resilience Scale (RS-13^81^) to assess resilience as a characteristic that moderates the negative effects of stress and promotes adaptation (sum score 0-91; α = .90).

### Visceral pain model and thresholding procedure

The study, with an overall duration of 2h, was performed at the University Hospital Essen. An overview of the experimental procedures is provided in Figure 4A-C. Initially, a thresholding procedure for visceral stimuli was accomplished using a well-established protocol^24,28,30–33^ to determine individual sensory, urge, and pain thresholds for rectal distensions used as an experimental model of visceral pain herein. To this end, an inflatable balloon attached to a pressure-controlled barostat system (modified ISOBAR 3 device; G & J Electronics, ON, Canada) was positioned 5 cm from the anal verge, allowing the application of graded rectal distension stimuli. This model simulates visceral sensations, including visceral pain, in a controlled and reproducible way and is widely used in human experimental research and clinical practice^82,83^.

**Figure 4.**
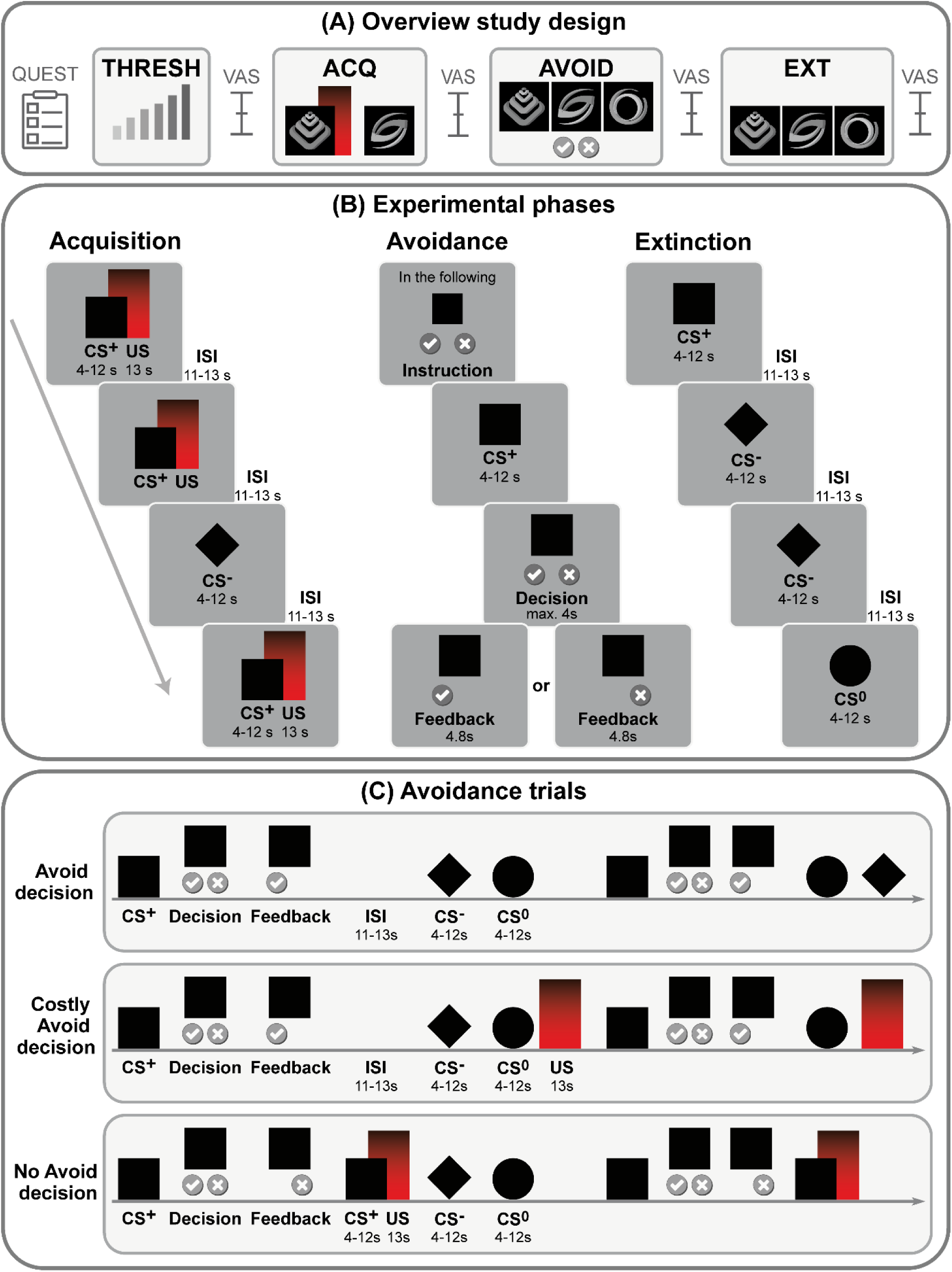
(A) Overview of procedures on the study day, (B) experimental acquisition (ACQ), avoidance (AVOID) and extinction (EXT) phases and (C) types of CS^+^ trials during the avoidance phase. Abbreviations: CS, conditioned stimulus; ISI, interstimulus interval; THRESH, thresholding procedure; US, unconditioned stimulus; VAS, visual analog scale.

To determine individual distension pressures for application of visceral pain stimuli during the following experimental phases, distensions of different perceptual intensities were applied in a double-random staircase procedure with random pressure increments of 2 - 6 mmHg, durations of 30 s, and a maximal distension pressure of 50 mmHg. After each stimulus, participants were prompted to rate the respective distension on a Likert-type scale with labels 1 = no perception, 2 = doubtful perception, 3 = urge to defecate, 4 = pain, not tolerable distension using a hand-held fiber optic response device (LUMItouch™, Photon Control Inc., Burnaby, BC, Canada).

The length of stimulation was chosen based on established methodology and to ensure a sufficiently long plateau phase of the respective stimulus intensity for the rating, during which pressures did not change. Sensory thresholds were defined as the pressure when ratings changed from 1 to 2, urge thresholds were defined as ratings of 3, and pain thresholds were determined as a change from 3 to 4. These individual thresholds were used as anchors to identify the intensity of a moderately painful but still tolerable US for repeated application during the subsequent experimental phases. Following established procedures for calibrating stimulus intensities based on their perceptual properties^84–87^, individual sensory thresholds were used to define the intensity of the unconditioned stimulus (US) as moderately painful but still tolerable. Specifically, participants first received a stimulus at the pressure corresponding to the highest rating of 3 (i.e., highest pressure inducing an urge to defecate). They rated the intensity of this stimulus on a 0-100 mm visual analogue scale (VAS), with endpoints labeled ‘not perceivable’ and ‘very painful.’ The pressure was then adjusted in 1 mmHg increments to identify a pressure just below the individual pain threshold. The final distension pressure was then applied as a US for 13 seconds and rated to ensure an adequately intense US targeted to elicit VAS ratings between 60 and 80 to induce a robust aversive response by the moderately painful stimulus intensity, while keeping the overall procedure during the experiment tolerable for repeated exposure in the subsequent experimental phases.

### Experimental paradigm

Based on a well-established protocol^28,30,32,33,36^ during the acquisition phase, two distinct abstract geometric shapes in various shades of gray, as depicted in Figure 4A were presented as conditioned stimuli (8 CS^+^, 8 CS^−^), and 8 visceral pain stimuli were delivered as US (Figure 4B). One visual cue (CS^+^) was contingently paired with the US in a 100% reinforcement schedule, and both stimuli co-terminated (differential delay paradigm). A second visual cue (CS^−^) was presented unpaired.

At the beginning of the subsequent avoidance phase, participants were instructed as follows: “In the following, you may be able to skip the pain stimulus when you see this symbol.”, while a CS^+^ was presented together with a checkmark (left) and a cross (right), representing the response buttons. They were prompted to indicate their decision either to avoid pain by pressing the left response button on the LUMItouch™ response device using their index finger or to subsequently receive pain by pressing the right button with their middle finger. During the avoidance phase (Figure 4C), 12 out of 16 CS^+^ trials involved an option to make a decision. Here, the CS^+^ was initially presented alone, then the two buttons to indicate the option to avoid or to not avoid appeared on the screen while the CS^+^ remained visible. When participants decided to receive pain (No Avoidance trials), the respective button remained on the screen for 4.8 seconds as feedback. Thereafter, CS^+^ was presented, and US was delivered as during the acquisition phase.

When participants decided to avoid the US, the corresponding button was displayed, but was not followed by the previously established contingent CS^+^-US pairing, but an interstimulus interval (ISI), initially deeming the decision to avoid as successful from the participants’ point of view. However, while in Avoidance trials, US remained absent until the next CS^+^ presentation, in Costly Avoidance trials, participants were confronted with an unsignaled US in a non-contingent manner and at a random time point following this decision and before the next CS^+^ trial, as previously accomplished^38^. These two types of trials were randomly distributed across the avoidance phase to prevent a reliable prediction of decision outcomes for the participant. This approach was chosen to overcome some previous limitations by experimentally modeling avoidance costs as the confrontation with an unexpected painful sensation concomitant with the loss of contingency. As fear avoidance is considered an active process, involving conscious and unconscious efforts to avoid situations perceived as threatening or painful, no response would have likely occurred due to inattentiveness and would have precluded a valid interpretation of any decision. Therefore, no response would have been classified as a No Avoidance decision.

As control conditions, four CS^+^ trials for which no decision could be made and 12 CS^−^ trials resembling those of the acquisition phase were implemented. In addition, a novel CS^0^ cue was introduced and presented in 12 trials as a control cue carrying no previously conditioned predictive value. During extinction (Figure 4B), CS^+^, CS^−^, and CS^0^ (8 trials each) were presented in the absence of US.

CS durations were jittered between 4 and 12 s to avoid conditioning to fixed time intervals, and US during acquisition and avoidance phases were delivered for 13 s. ISI during all experimental phases had a variable duration between 11 and 13 seconds. Overall durations were 9:11 min for the acquisition phase, between 16:17 min and 17:15 min for the avoidance phase (depending on individual decision patterns), and 10:31 min for the extinction phase. All stimuli were presented in a balanced and pseudo-randomized order using Presentation® software (Neurobehavioral Systems, Albany, CA, USA) with the following constraints: no more than two consecutive repetitions of the same CS or trial type and equal numbers of each CS.

### Behavioral and self-report measures

Before (baseline) and after each experimental phase, measures of CS valence, fear of CS, US intensity, and fear of US were assessed using a hand-held LUMItouch™ device, as previously accomplished^24,31^. For CS valence, the respective CS was presented contemporaneously for 10 seconds, and participants were asked to respond to the question “How do you perceive this stimulus?” on a visual analog scale (VAS) with endpoints labeled “very pleasant” (-100) and “very unpleasant” (+100) with neutral perception indicated in the middle of the scale (0). To ensure that all participants had acquired pain-related fear as an essential prerequisite for examining subsequent avoidance behavior and extinction, successful differential fear acquisition was defined as an inclusion criterion for further analyses, based on previous literature^88^. Valence ratings were used to assess this as indicators of learned emotional aversion toward the CS^+^. In an initial blinded analysis, one participant’s ratings indicated a lack of fear acquisition, as the CS^+^ was rated as more pleasant following acquisition compared to baseline. This participant was therefore excluded from subsequent analyses.

US intensity was assessed using the question “How intense was the distension stimulus?” with endpoints labelled “unperceivable” (0) and “extremely painful” (100) directly following the application of a US. The question and endpoints encompassing both sensory intensity and painfulness were chosen to reflect participants’ subjective perception of stimulus strength across the full range of visceral sensations, which is comparable to the well-established thresholding procedure, capturing first sensation, unpleasant urgency, and painfulness as dimensions of visceral perception of increasing intensity.

Fear of CS and US, respectively, was determined by asking participants, “How fear-inducing is this symbol / was the distension stimulus?” with 0 indicating “not at all” and 100 indicating “very much”. Additionally, CS-US contingency awareness was evaluated following acquisition, avoidance, and extinction phases by presenting CS^+^, CS^−^ and CS^0^ and prompting participants to respond to the question “How often was this symbol followed by a distension stimulus?” on a VAS with endpoints labeled “never” (0) and “always” (100).

To distinctly contemplate individual decision patterns during the avoidance phase, trial-by-trial decisions to avoid or receive pain were assessed and reaction times for each decision (measured in ms and transformed to s) were logged using Presentation® software.

### Statistical analyses

A total of N = 5 datasets were excluded (technical error resulting in a loss of experimental data, N = 1; lack of fear learning during acquisition, N = 1; structural brain abnormality, N = 1; premature termination of the experiment, N = 2), resulting in a final sample of N = 33 participants (15 women, 18 men) included in statistical analyses.

Analyses were conducted using IBM SPSS Statistics 27.0 (IBM Corporation, Armonk, NY, USA). Initially, normal distribution of outcomes was confirmed using Kolmogorov–Smirnov tests, and two-sided parametric testing was subsequently applied. Putative sex differences in relevant psychological measures and thresholds as proxies of visceral sensitivity were excluded using two-sample t-tests. This was done in light of previous and partly conflicting findings in the field of pain, reporting on sex differences in pain perception for some modalities and measures, yet no differences between healthy men and women in others^89^, including visceral pain sensitivity^90^.

Repeated measures analysis of variance (RM-ANOVA) were performed for US intensity, US valence and for fear of US ratings with the within-subject factor *time* (baseline, acquisition, avoidance) to evaluate effects of pain-related learning and avoidance on different facets of pain perception. Exploratory correlational analyses were conducted to assess possible associations of US-related measures with avoidance decisions.

For ratings of CS-US contingencies, RM-ANOVA including the within-subject factors *stimulus type* (CS**^+^**, CS**^-^**, CS^0^) and time (acquisition, extinction) followed by post hoc paired t-tests, was calculated.

RM-ANOVA were further conducted for CS valence and for fear of CS, including the within-subject factors *stimulus type* (CS**^+^**, CS**^-^**, CS^0^) and *time* (baseline, acquisition, avoidance, extinction). Complementing and extending this approach, hierarchical stepwise regression analyses evaluating to what extent pain-predictive cue properties could explain variance in self-reported correlates of CS^+^ fear and valence were further calculated. The inclusion of variables was based on a *p*-value of *F* ≤ .050, and exclusion on the probability of *F* ≥ .100, and multicollinearity was initially inspected, resulting in VIF scores near 1, indicating low collinearity. CS^+^ valence and fear ratings from previous experimental phases, as well as the percentage of avoidance decisions, were entered in a first stage. Trait anxiety, chronic stress, and coping together with uncertainty intolerance were added in a second stage to explore whether trait psychological characteristics relevant to avoidance could further improve model fit.

To scrutinize avoidance behaviors, the percentage of avoidance decisions was considered and individual trial-by-trial decisions as well as reaction times per decision were analyzed using generalized linear mixed models for fixed effects of *trial* and *prior experience* (unsignaled US, signaled US, avoided US) to account for inter-as well as intraindividual variability and Fisher’s Exact Tests were conducted to compare proportions of the respective decisions. Adding a random intercept for each participant and allowing variation for the factors *trial*, *prior experience* and *participants* by adding random slopes for these factors did not further improve model fit for analyzing trial-by-trial decisions (Bayes = 1635.33). Of note, due to a technical error, single decisions were not available for some participants, resulting in missing data for together 26 out of 363 decisions (∼7%, randomly distributed across participants and trials) during the avoidance phase. These missing data were not interpolated or deleted, as the primary focus was on investigating avoidance behavior based on individual prior decisions and experiences.

Exploratory correlational analyses were conducted to identify putative associations of avoidance decision patterns with psychological trait measures.

All ANOVA results are reported with Greenhouse-Geisser correction to account for a possible violation of the sphericity assumption, and results of post hoc paired t-tests were Bonferroni-corrected for multiple comparisons. The alpha level for accepting statistical significance was set at *p* < .05. Data are shown as mean ± standard error of the mean (SEM), unless indicated otherwise, and effect sizes are provided as *η_p_^2^*or Cohen’s *d,* respectively.

## Acknowledgements

This work was supported by the Deutsche Forschungsgemeinschaft (DFG, German Research Foundation) project number 316803389 – SFB 1280, Project A10. The funding agency had no role in the conception, analysis, or interpretation of the data.

We would like to thank Miller Haimour for support in data acquisition. We further thank Dr. Katarina Forkmann and Dr. Robert Pawlik for excellent technical support.

## Author contribution statement

F.L. and A.I. contributed to the conception of the work and secured funding. F.L., A.K., and A.I. acquired and analyzed the data. F.L., S.E., and A.I. interpreted the findings, drafted, and revised the work. All authors reviewed and approved the final manuscript.

## Data availability

The data that support the findings of this study are not openly available but are available from the corresponding author upon reasonable request. Data are located in controlled access data storage at Ruhr University Bochum.

## Additional information

The authors declare that they have no known competing financial interests or personal relationships that could have appeared to influence the work reported in this paper.

## Notes

### Competing Interest Statement

The authors have declared no competing interest.

